# Prediction of the virus incubation period for COVID-19 and future outbreaks

**DOI:** 10.1101/2020.05.19.104513

**Authors:** Ayal B. Gussow, Noam Auslander, Yuri I. Wolf, Eugene V. Koonin

**Author notes:** These authors contributed equally (Ayal B. Gussow, Noam Auslander).

## Abstract

A crucial factor in mitigating respiratory viral outbreaks is early determination of the duration of the incubation period and, accordingly, the required quarantine time for potentially exposed individuals. Here, we explore different genomic features of RNA viruses that correlate with the incubation times and provide a predictive model that accurately estimates the upper limit incubation time for diverse viruses including SARS-CoV-2, and thus, could help control future outbreaks.

## Main Text

The recent outbreak of the novel SARS-CoV-2 coronavirus and the resulting COVID-19 disease has led to an unprecedented worldwide emergency^1^. Per World Health Organization (WHO) recommendations, numerous countries have taken severe preventive measures to combat and stem the spread of the virus. A key effective measure recommended by the WHO in viral outbreaks is enforcing a period of quarantine on individuals that are suspected to have come in contact with the causative agent until they are proven clean of infection^2,3^. The length of the quarantine typically depends on the time from virus exposure to the emergence of symptoms, i.e. the incubation period. The duration of the incubation period is specific to the causative virus^4^. Underestimation of the incubation time could lead to infected individuals being prematurely released from quarantine and spreading the disease, whereas overestimation can have a debilitating economic impact and cause detrimental psychological effects^5^. Therefore, knowledge of the upper limit of a virus incubation period is crucial to effectively combat and prevent outbreaks while minimizing the negative consequences of the quarantine.

The length of the incubation period varies both across and within virus families^4^. To our knowledge, genomic features (if any) that correlate with the incubation time are currently unknown. There is therefore a vital need for methods to predict the incubation periods of emerging viruses. Such methods can be deployed in future viral outbreaks for early, accurate inference of the incubation period and immediate implementation of optimized quarantining interventions that will mitigate the spread of the virus while minimizing the negative societal impact^6^.

Here, we analyzed different genomic characteristics of respiratory, non-segmented, single-strand RNA (ssRNA) viruses and identified features that are predictive of the incubation time, and are generalizable across virus families. Based on these features, we developed an elastic net regression model that predicts virus incubation periods. We extensively validated the robustness of this model and the selected features for the prediction of the incubation period across diverse viruses and virus families, to enable accurate early estimation of the incubation period for future outbreaks.

We focused on non-segmented ssRNA viruses that cause respiratory infections, to minimize confounders to the incubation period that could result from different genome structures and/or different infected tissues and cell types. In total, we collected information for 14 viruses from 4 families. Given that the quarantine time is defined as the upper limit of virus incubation time, we extracted the upper estimates of the incubation periods for all viruses in the analyzed set (Supp. Table 1, see Methods for details). We then curated 8 features that we hypothesized could be relevant for the incubation period (Figure 1a), based on the complete genome nucleotide sequences and within-population genome alignments of all sequenced strains of each virus (Supp. Datasets 1 and 2, see Methods for details). Analysis of the pairwise associations between these features (Fig. 1a) confirmed some previously reported associations, such as the negative correlation between genome length and mutation rate^7^ and the positive correlation between GC content and codon adaptation index^8^ (CAI) (Fig. 1b). Strikingly, our findings suggest that the mutation rate of SARS-CoV-2 is substantially lower than that of other human coronaviruses (CoV), including its closest human-infecting relative, SARS-CoV (with an average of 1.4e-3 and 7.8e-5 transitions per branch point per nucleotide for SARS-CoV and SARS-CoV-2, respectively, Fig. 1b, see Methods for details). Preliminary reports on SARS-CoV-2 genome evolution indicate a similar trend^9^.

**Figure 1.**
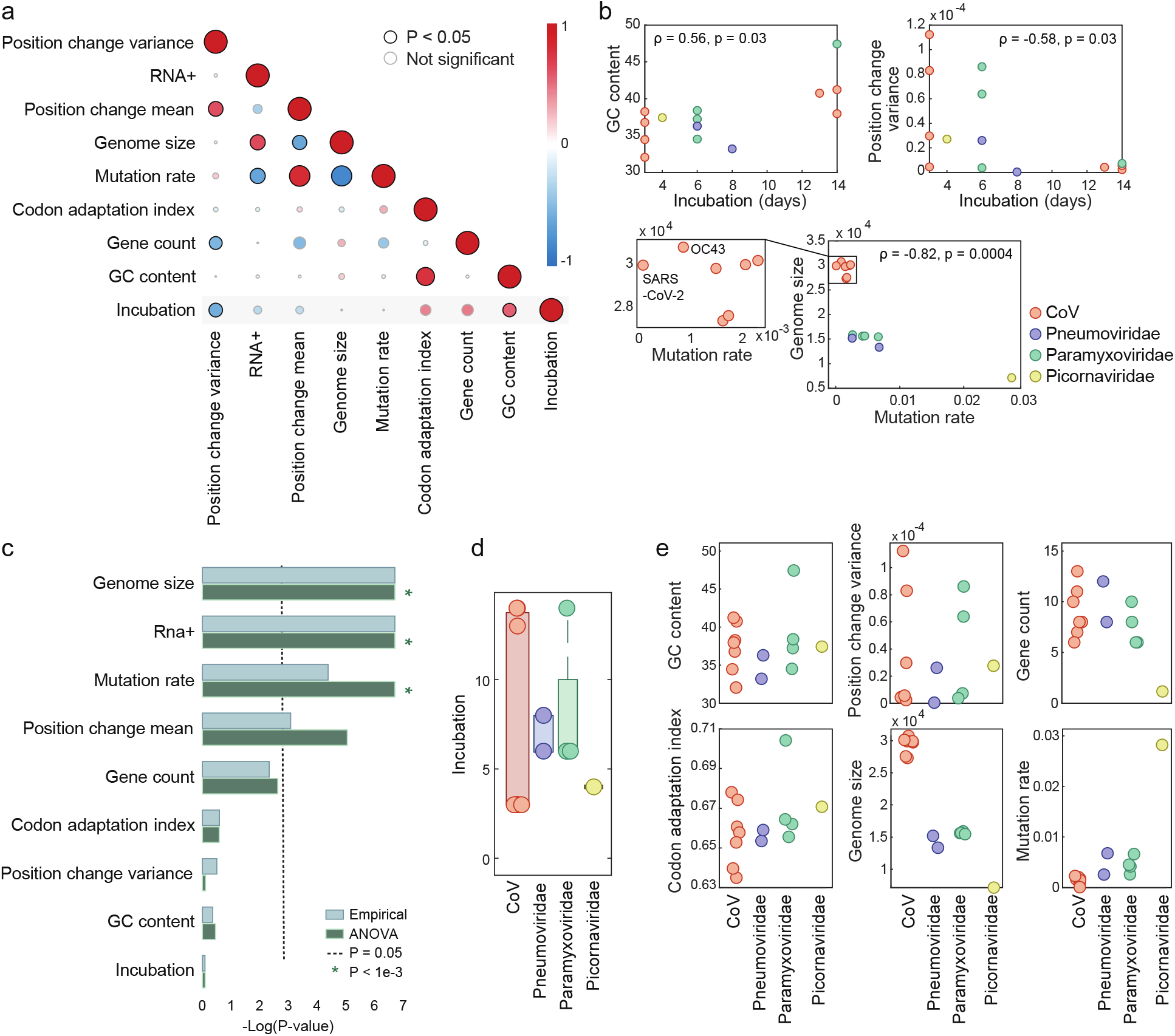
Genomic features of ssRNA viruses causing respiratory infections. **(a)** Pairwise correlation matrix across all features. A description of feature construction is given in the Methods section. Each circle indicates Spearman’s ρ between two features. The colors represent the rank-correlation coefficients (red indicates positive correlation and blue indicates negative correlation), and the circle sizes correspond to significance (p-value), where significant correlations (p-value <0.05) are circled in black. **(b)** Scatter plots illustrating the relationships between features across four virus families. **(c)** Estimation of the features association with the virus family, based on p-values (-log scaled) from two tests applied (see Methods for details). The cutoff (p-value = 0.05) is indicated with a dashed line. Lower values correspond to features that are not significantly associated with a virus family. **(d)** Boxplot and overlaid dot plot of the incubation periods across viral families. **(e)** Dot plots of different features across virus families. The features shown in the upper panels are family-generic, and those in the bottom panels are family-specific.

We then sought to select features to be used for a predictive model of the incubation time. To avoid confounding the model with features that are primarily driven by virus family, we formally quantified whether a given feature is significantly associated with the family identity. To this end, we applied two complementary approaches, namely, analysis of variance (ANOVA) and an empirical, non-parametric test, to estimate, for each feature, whether it varies more across virus families than within each family (see Methods for details). The results obtained with the two approaches were equivalent, demonstrating that half of the considered features vary more between families than within families, and therefore may confound the model (Fig. 1c). We denote such features family-specific. By contrast, we found that the incubation time was not significantly associated with virus family (Fig. 1d), supporting selection of features that are not family-specific to train a model; we denote such features family-generic. Four other features were found to be family-generic: GC content of the virus genome; variance of the number of different nucleotides observed per position in the alignment of the virus strains; number of protein-coding genes in the virus genome; codon adaptation index (CAI) (Fig. 1e). Thus, these four features were included in the model.

Next, we sought to divide the analyzed dataset into training and test sets. To maintain a large enough independent test set, we trained an elastic net model on the 7 human-infecting viruses of the family *Coronaviridae* (Fig. 2a), using the four family-generic features. We found that this model, which was trained on a single viral family, generalized well to viruses from the three other families (Fig. 2b). The test mean absolute error was 1.63 days (Fig. 2b), attesting to a close estimation of the upper limit of the incubation time in an independent data set. Moreover, the model predictions strongly correlated with the ranks of the assigned incubation periods in the test set (Spearman’s ρ = 0.91, p-value = 0.005). Specifically, for the virus with the longest known incubation period, measles, the longest incubation time, 9.7 days, was predicted. Although measles was assigned an upper limit incubation period of 14 days in our data, the majority of the available reports are indeed in the range of 9-12 days^10^. The second longest incubation period was also correctly assigned, to respiratory syncytial virus (RSV), with a prediction of 9.1 days, compared to an assigned period of 8 days in our data. For parainfluenza 1-3, the model predicted 7.3, 4.9, and 6.2 days, respectively, closely approximating the assigned 6 days. Metapneumovirus was similarly accurately predicted to have a 6.5 days incubation period, within half a day of its assigned 6 days. Finally, the shortest incubation time predicted was correctly assigned to rhinovirus, with a prediction of a 1.2 day incubation period. Although rhinovirus was assigned a 4 day incubation period in our data, most of the cases show symptoms within one day^11^.

**Figure 2.**
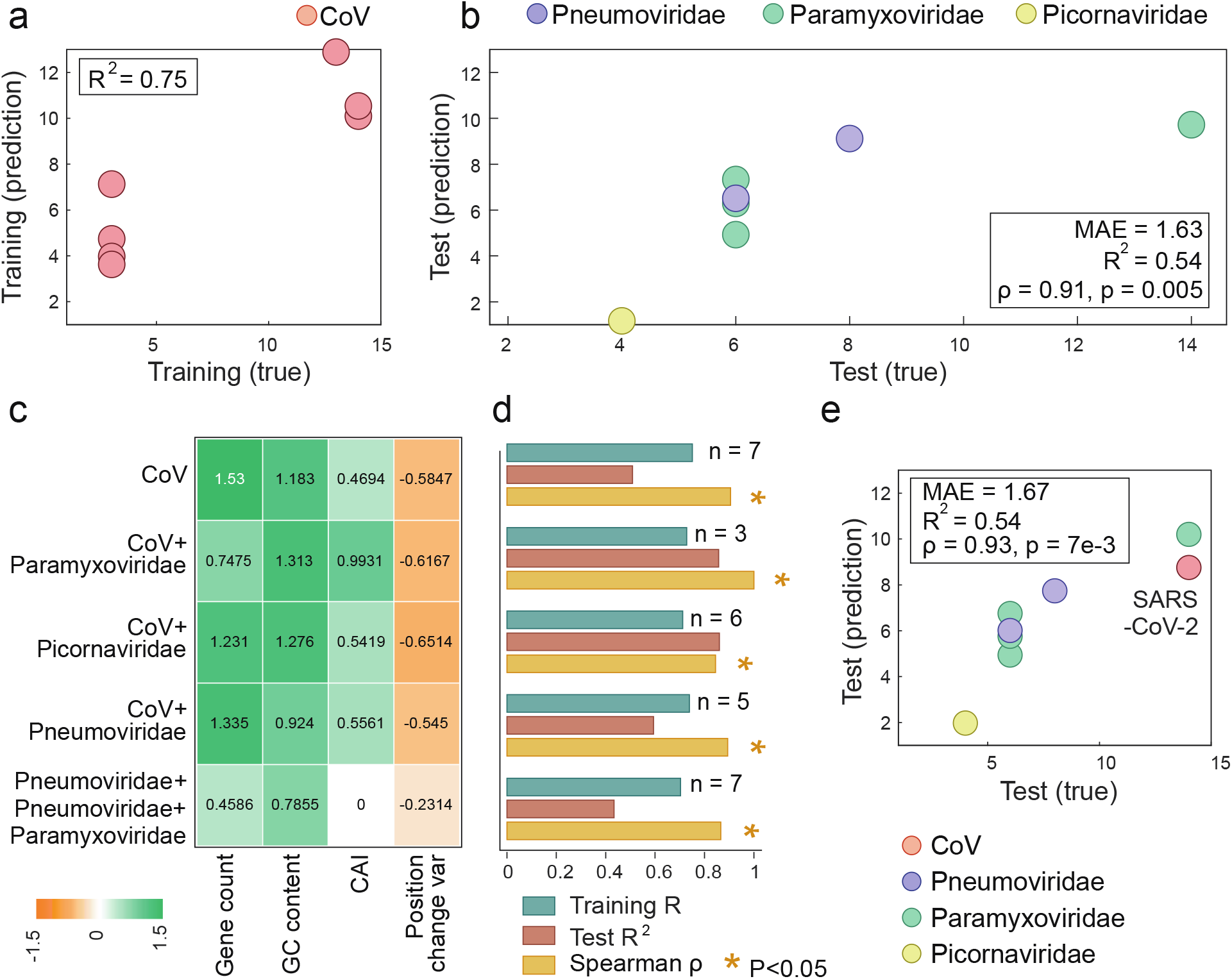
Assessment of the elastic net model for virus incubation time prediction. **(a)** A scatter plot of the incubation periods of the CoV training set compared to the model predictions, with the model R^2^ in the upper left corner. **(b)** A scatter plot of the incubation periods of the test set compared to the model predictions. The box in the bottom right corner contains: Spearman’s ρ between the predictions and the true values; the P-value of the Spearman’s ρ; the model R^2^; the mean absolute error (MAE). **(c)** A heatmap of the coefficients of each feature using different training sets. **(d)** A bar plot of the model performance metrics using different subsets of training and testing data, with the number of samples in the testing data for each subset indicated. **(e)** A scatter plot of the incubation periods of the test set compared to the model predictions when SARS-CoV-2 is left out of training. The box in the top left corner shows the model’s performance metrics.

Exploration of the model indicated that the strongest predictive features were the number of protein-coding genes and GC content, with higher values in either feature corresponding to a longer incubation time (Fig. 2c). Elucidation of the mechanisms behind these associations will require extensive experimental work. A straightforward, even if, likely, over-simplified explanation could be that the larger number of genes to be translated by the virus lengthens its replication cycle, under the assumption that the number of translation initiation events and/or subgenomic RNAs that need to be transcribed are rate-limiting factors in virus reproduction. Similarly, a higher GC content leads to the formation of stable secondary structures in the virus RNA, with higher kinetic barriers that the ribosome then needs to disrupt during translation, resulting in longer translation times^12^. Thus, one possible explanation for the association between the number of protein-coding genes and the GC content and longer incubation periods is that the longer cumulative translation time extends the replication cycle and consequently, the incubation period. Alternatively or additionally, extra genes could contribute to more complex interactions of the virus with the host organism, resulting in a longer incubation times. In particular, the highly virulent coronaviruses with long characteristic incubation periods are known to encode additional, accessory proteins compared to low virulence viruses that have shorter incubation times^13,14^. The accessory genes are dispensable for virus reproduction in cell culture and have been implicated in virus-host interactions^15^. Recent preliminary analysis has shown that some of these additional genes encode proteins containing distinct immunoglobulin-like domains, which is compatible with roles in interactions with the immune system of the host^16^.

To assess the robustness of the selected features, we tested models trained with different partitioning of the data into train and test sets. We found that these changes did not significantly change the performance of the models, further attesting to the robustness of the signal obtained using the four family-generic features (Fig. 2d). By contrast, a model trained with family-specific features does not generalize to the test set, and one trained using a mixture of family-generic and family-specific features disregards the latter by nullifying their coefficients (Supp. Fig. 1), further demonstrating the efficacy of relying on family-generic features only. The coefficients assigned to the family-generic features did not vary substantially across different training sets, confirming that the method is not particularly sensitive to the data used for training (Fig. 2c). Nevertheless, the high performance of the model that is trained exclusively on CoV seems to suggest that this virus family provides a good representation of the incubation period, and/or that training on a single family is preferable given the small dataset and the possibility of confounding effects.

To evaluate the utility of our model, we examined how this method could have performed during the early stages of the current COVID-19 pandemic. To this end, we removed SARS-CoV-2 from the training data and trained the model on the remaining 6 CoV only, with the caveat that this training set is poorly balanced as it contains only 2 viruses with incubation times longer than 3 days, and therefore, might underestimate when predicting viruses with long incubation times. The incubation period of SARS-CoV-2 is still being determined, with the recommended quarantine time conservatively set at 14 days although recent reports indicate that 97.5% of patients who will develop symptoms do so within a 95% confidence interval of 8.2-15.6 days^17^. Despite having been trained on an imbalanced training set, the model predicts an incubation period of 8.8 days for SARS-CoV-2, correctly placing SARS-CoV-2 in the upper range of incubation periods, well within the confidence interval, and predicting an incubation period duration during which the vast majority of symptomatic patients will have shown symptoms^17^. Clearly, this estimate could have been useful in mitigating the COVID-19 pandemic (Fig. 2e; similar analysis for the other CoV is provided in Supp. Fig. 2).

In summary, we examined here genomic features that could be predictive of the virus incubation times of human pathogenic ssRNA viruses, some of which are major causative agents of respiratory pandemics^18^. We identified four family-generic genomic features that consistently predict the incubation periods with high accuracy. Using these features, we developed a robust model that is predictive of incubation times and can directly facilitate early and accurate estimation of the required quarantine time for future pandemics.

## Methods

### Sequence Datasets

Reference genome sequences and GenBank files were downloaded from the NCBI^19^ for each virus (Supp. Table 1, Supp. File 1). For each virus, additional strains were downloaded from the NCBI and aligned using Mafft^20^ v7.407 with default parameters, resulting in an alignment file for each of the 14 viruses (Supp. File 2). Phylogenetic trees were generated for each virus based on the alignment using FastTree^21,22^ with the “-nt” parameter.

### Incubation period assignment

The incubation time for each of the 14 viruses was collected from literature (Supp. Table 1). As incubation periods vary, where possible, the upper limit was used, and a consensus of reports was followed. We note that there are small variations in reports of the incubation times, and the assigned values represent the best approximation as explained below. Changing the assigned incubation times within the range of reports maintains a similarly high performance of the trained model (Supp. Fig. 2).

The rationale for the selected incubation times was as follows:

a. *229E-CoV, HKU1-CoV, NL63-CoV and OC43-CoV*. For these coronaviruses, which are causative agents of common colds, a 3 day incubation period was assigned, following the majority of reports^23,24^.
b. *MERS-CoV*. A 14 day incubation period was assigned to MERS-CoV, per previous reports^25^.
c. *SARS-CoV.* The estimates show 13 days as an upper limit in the majority of reports^4,26^.
d. *SARS-CoV-2*. Although more data is needed to assess the incubation period of SARS-CoV-2, we set the duration to 14 days given the recommended quarantine times^2^.
e. *Measles virus*. There is a considerable range of reported incubation periods, with the majority of reports indicating 9-12 days and some reports going several days beyond that. Thus, we set the incubation period to 14 days^10^.
f. *Respiratory syncytial virus (RSV).* For RSV, the incubation period was set to 8 days, in accordance with the higher range of the majority of reports^27^.
g. *Parainfluenza*. The parainfluenza incubation period was consistently reported to be between 2-6 days, and therefore was assigned the upper limit of 6 days^4^.
h. *Rhinovirus*. Per previous reports, a 4 day incubation period was assigned to Rhinovirus^4^.
i. *Metapneumovirus*. 6 days was assigned to human metapneumovirus, given the commonly reported range of 4-6 days^28^.

### Features

Genomic features were calculated as follows:

a. *Genome length*. The number of nucleotides in the reference genome sequence.
b. *Number of genes*. The number of genes in the reference genome’s associated GenBank file. We verified for each virus that there were no undetected genes within its genome using MetaGeneMark^29^ gene prediction software.
c. *Positive or negative strand RNA.* Whether the RNA virus is positive strand or negative strand. This was set to 1 if the virus has a positive strand genome, and to 0 if it has a negative strand genome.
d. *Codon adaptation index (CAI)*. The CAI was used to analyze the codon usage bias of each virus in comparison to human. The CAI was calculated by concatenating all the coding sequences (CDS) in each virus reference genome GenBank file and using the Biopython^30,31^ software package implementation with a reference human codon usage table^32^.
e. *GC content*. This was calculated for each reference genome using Biopython^30^.
f. *Mutation rate.* Raw mutation rates were estimated per each virus genome alignment, without accounting for selection bias, by detecting the ancestral base for every base in the genome for every non-leaf node in the tree using maximum parsimony. Then, at each branch point, the transitions between both sides of the branch were counted, and the average count was then divided by the length of the genome for the final estimate.
g. *Average and variance of the changes in each position of the alignment.* The change in alignment position is defined as the number of different values observed in each position of the virus alignment, divided by the number of strains in the alignment. The average and variance of these values are used as features.

### Evaluation of the specificity of the features for virus families

We sought to evaluate, for each feature, whether it is associated with the identity of the virus family which would be a potential confounder to the model. We hence searched for features whose variance within each virus family was not significantly smaller than its overall variance. To this end, each feature was evaluated using two methods.

The first method is a one-way Analysis of variance (ANOVA). One-way ANOVA tests the null hypothesis that the means of the measurement variable are the same for the different categories of data, against the alternative hypothesis that they are not all the same. Hence, lower assigned p-values signify that the null hypothesis is rejected, and that different viral families have different population means with respect to each feature. We therefore consider features assigned with a P-value greater than 0.05, for which we could accept the null hypothesis, and could not conclude that the feature mean was associated with the viral family. The ANOVA test was implemented in Python using the f_oneway function in the SciPy^33^ package.

The ANOVA test assumes that the samples are independent, taken from normally distributed populations with equal standard deviations between the groups. These assumptions, that must be satisfied for the associated p-value to be valid, are not guaranteed and are difficult to evaluate. We hence implemented a second, empirical test, which is not parametric and does not rely on any assumptions. This empirical test evaluates, for a given feature, if its variance within virus families is smaller than would be observed by random assignment of families to viruses. We reason that a feature which is associated with the virus family would have significantly smaller variance within the true family assignment than within a random family assignment. The null hypothesis is that the variance of the features within each family is similar to the variance across families, and the alternative hypothesis is that the variance of the features within each family is smaller than the variance across families. To perform the empirical test, the feature variance within each virus family is calculated and averaged. Next, the feature values are randomly permuted 1,000 times and the same calculation is performed, to generate a null distribution. Let the number of times the variance of the permuted values is less than the variance of the real values be *X*. The p-value is calculated as (*X* + 1)/(1000 + 1). Thus, a lower p-value indicates that the feature’s within-family variance is smaller than our null expectation. We search for features with a p-value greater than 0.05, for which we conclude that the variance within the actual families is not smaller than that within randomly assigned families. This evaluation does not necessarily indicate that the family-specific features are poor predictors, rather, that, with the data available, it would not be possible to discern whether the signal from these features is primarily driven by the variation between virus families.

### Elastic Net Model

The elastic net method^34^ is a generalization of LASSO using Ridge regression shrinkage, where the naïve estimator 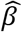 is a minimizer of the criterion *L*(λ_1_, λ_2_, β) by:

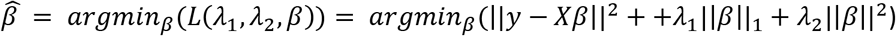

for any fixed, non-negative λ_1_, λ_2_. Elastic net was chosen because it has characteristics of both LASSO and Ridge regression, which are controlled by the penalties coefficients, thus outperforming other regularization and variable selection approaches^34^.

The elastic net model was constructed in Python using the scikit-learn^35^ ElasticNet function with default parameters. The features were standardized before training, with the same standardization parameters used in training applied to test data before prediction.

## Data availability

Virus genome sequence alignments used for this analysis are provided as supplementary information.

## Code availability

Our model, features and the accompanying code are available at https://github.com/noamaus/incubation-model.

## Acknowledgements

This research was supported by the Intramural Research Program of the National Library of Medicine at the NIH.

## Notes

### Competing Interest Statement

The authors have declared no competing interest.

## References

1. World Health Organization. Statement on the second meeting of the International Health Regulations (2005) Emergency Committee regarding the outbreak of novel coronavirus (2019-nCoV). https://www.who.int/news-room/detail/30-01-2020-statement-on-the-second-meeting-of-the-international-health-regulations-(2005)-emergency-committee-regarding-the-outbreak-of-novel-coronavirus-(2019-ncov) (2020).

2. World Health Organization. Considerations for quarantine of individuals in the context of containment for coronavirus disease (COVID-19). https://www.who.int/publications-detail/considerations-for-quarantine-of-individuals-in-the-context-of-containment-for-coronavirus-disease-(covid-19) (2020).

3. Gupta, A. G., Moyer, C. A. & Stern, D. T. The economic impact of quarantine: SARS in Toronto as a case study. J. Infect. 50, 386–93 (2005).

4. Lessler, J. et al. Incubation periods of acute respiratory viral infections: a systematic review. Lancet Infect. Dis. 9, 291–300 (2009).

5. Hawryluck, L. et al. SARS control and psychological effects of quarantine, Toronto, Canada. Emerg. Infect. Dis. 10, 1206–12 (2004).

6. Koo, J. R. et al. Interventions to mitigate early spread of SARS-CoV-2 in Singapore: a modelling study. Lancet. Infect. Dis. (2020) doi:10.1016/S1473-3099(20)30162-6.

7. Sanjuán, R., Nebot, M. R., Chirico, N., Mansky, L. M. & Belshaw, R. Viral Mutation Rates. J. Virol. 84, 9733–9748 (2010).

8. Knight, R. D., Freeland, S. J. & Landweber, L. F. A simple model based on mutation and selection explains trends in codon and amino-acid usage and GC composition within and across genomes. Genome Biol. 2, RESEARCH0010 (2001).

9. Jia, Y. et al. Analysis of the mutation dynamics of SARS-CoV-2 reveals the spread history and emergence of RBD mutant with lower ACE2 binding affinity. bioRxiv 2020.04.09.034942 (2020) doi:10.1101/2020.04.09.034942.

10. Naim, H. Y. Measles virus. Hum. Vaccin. Immunother. 11, 21–26 (2015).

11. Harris, J. M. & Gwaltney, J. M. Incubation Periods of Experimental Rhinovirus Infection and Illness. Clin. Infect. Dis. 23, 1287–1290 (1996).

12. Qu, X. et al. The ribosome uses two active mechanisms to unwind messenger RNA during translation. Nature 475, 118–21 (2011).

13. Liu, D. X., Fung, T. S., Chong, K. K.-L., Shukla, A. & Hilgenfeld, R. Accessory proteins of SARS-CoV and other coronaviruses. Antiviral Res. 109, 97–109 (2014).

14. Cui, J., Li, F. & Shi, Z.-L. Origin and evolution of pathogenic coronaviruses. Nat. Rev. Microbiol. 17, 181–192 (2019).

15. Menachery, V. D. et al. MERS-CoV Accessory ORFs Play Key Role for Infection and Pathogenesis. MBio 8, (2017).

16. Tan, Y., Schneider, T., Leong, M., Aravind, L. & Zhang, D. Novel Immunoglobulin Domain Proteins Provide Insights into Evolution and Pathogenesis Mechanisms of SARS-Related Coronaviruses. bioRxiv 2020.03.04.977736 (2020) doi:10.1101/2020.03.04.977736.

17. Lauer, S. A. et al. The Incubation Period of Coronavirus Disease 2019 (COVID-19) From Publicly Reported Confirmed Cases: Estimation and Application. Ann. Intern. Med. 172, 577 (2020).

18. Carrasco-Hernandez, R., Jácome, R., López Vidal, Y. & Ponce de León, S. Are RNA Viruses Candidate Agents for the Next Global Pandemic? A Review. ILAR J. 58, 343–358 (2017).

19. Agarwala, R. et al. Database resources of the National Center for Biotechnology Information. Nucleic Acids Res. 46, D8–D13 (2018).

20. Katoh, K. & Standley, D. M. MAFFT multiple sequence alignment software version 7: improvements in performance and usability. Mol. Biol. Evol. 30, 772–80 (2013).

21. Price, M. N., Dehal, P. S. & Arkin, A. P. FastTree 2 – Approximately Maximum-Likelihood Trees for Large Alignments. PLoS One 5, e9490 (2010).

22. Price, M. N., Dehal, P. S. & Arkin, A. P. Fasttree: Computing large minimum evolution trees with profiles instead of a distance matrix. Mol. Biol. Evol. 26, 1641–1650 (2009).

23. Bradburne, A. F., Bynoe, M. L. & Tyrrell, D. A. Effects of a ‘new’ human respiratory virus in volunteers. BMJ 3, 767–769 (1967).

24. Tyrrell, D. A. J., Cohen, S. & Schilarb, J. E. Signs and symptoms in common colds. Epidemiol. Infect. 111, 143–156 (1993).

25. Nishiura, H., Miyamatsu, Y. & Mizumoto, K. Objective Determination of End of MERS Outbreak, South Korea, 2015. Emerg. Infect. Dis. 22, 146–148 (2016).

26. Meltzer, M. I. Multiple Contact Dates and SARS Incubation Periods. Emerg. Infect. Dis. 10, 207–209 (2004).

27. Madge, P., Paton, J. Y. Y., McColl, J. H. H. & Mackie, P. L. K. L. Prospective controlled study of four infection-control procedures to prevent nosocomial infection with respiratory syncytial virus. Lancet 340, 1079–1083 (1992).

28. Schweon, S. J. Human metapneumovirus. Nursing (Lond). 43, 62–63 (2013).

29. Zhu, W., Lomsadze, A. & Borodovsky, M. Ab initio gene identification in metagenomic sequences. Nucleic Acids Res. 38, e132–e132 (2010).

30. Cock, P. J. A. et al. Biopython: freely available Python tools for computational molecular biology and bioinformatics. Bioinformatics 25, 1422–3 (2009).

31. Sharp, P. M. & Li, W.-H. The codon adaptation index-a measure of directional synonymous codon usage bias, and its potential applications. Nucleic Acids Res. 15, 1281–1295 (1987).

32. Nakamura, Y. Codon usage tabulated from international DNA sequence databases: status for the year 2000. Nucleic Acids Res. 28, 292–292 (2000).

33. Virtanen, P. et al. SciPy 1.0: fundamental algorithms for scientific computing in Python. Nat. Methods 17, 261–272 (2020).

34. Zou, H. & Hastie, T. Regularization and variable selection via the elastic net. J. R. Stat. Soc. Ser. B (statistical Methodol. 67, 301–320 (2005).

35. Pedregosa, F. et al. Scikit-learn: Machine Learning in Python. J. Mach. Learn. Res. 12, 2825–2830 (2011).

