## Supplementary material for "Prediction of the virus incubation period for COVID-19 and future outbreaks": Supp. Table 1

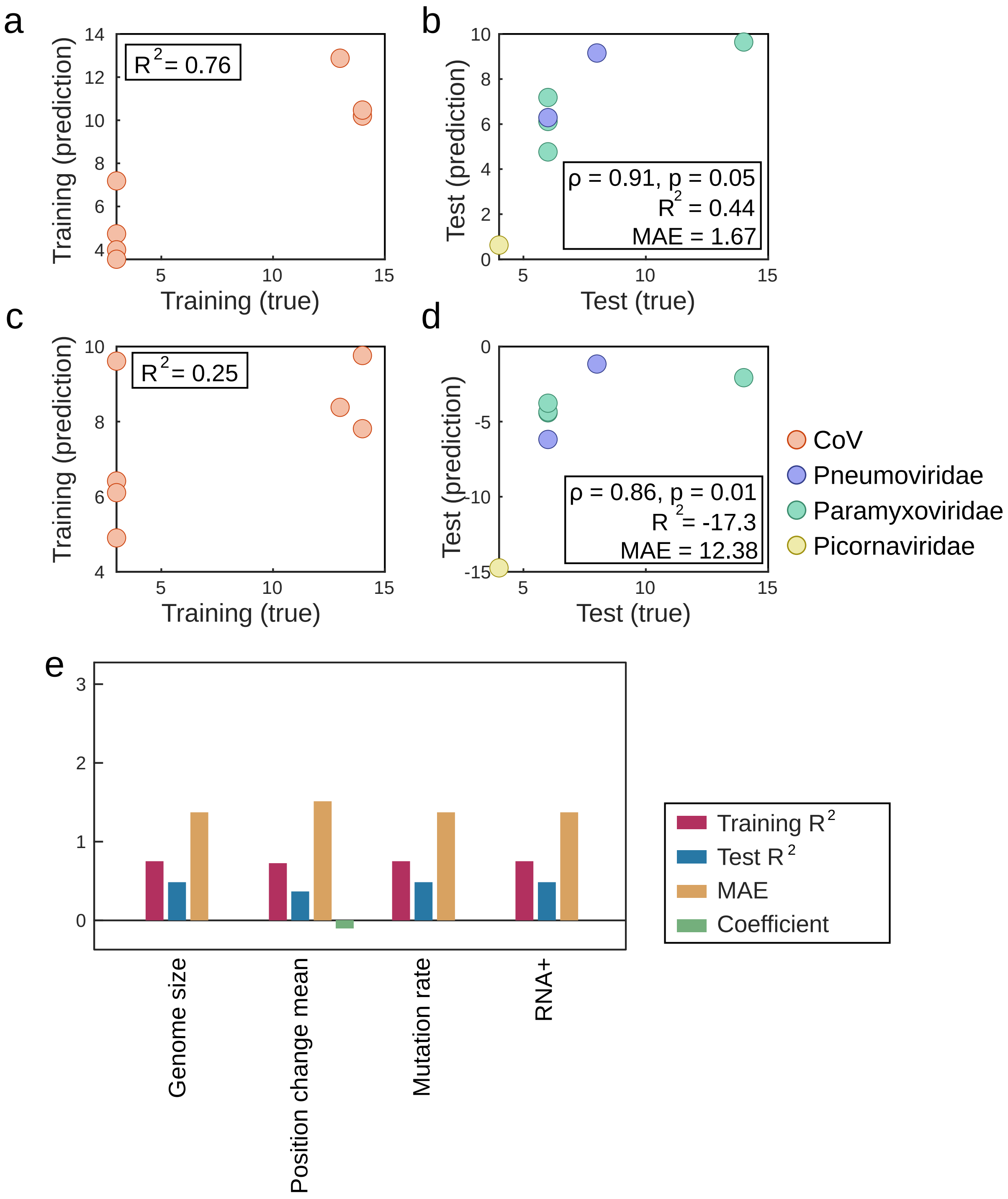


**Supplementary Figure 1. Performance of models trained with family-generic features.**

- **(a)** A scatter plot of the incubation periods of the CoV training set compared to the model predictions when trained using all features (family-generic and family-specific features). The training $R^2$ is slightly improved compared to the family-generic model in Figure 2a.

**(b)** A scatter plot of the incubation periods of the test set compared to the model predictions, when trained using all features (family-generic and family-specific features). The box in the bottom right corner contains: Spearman's ρ between predictions and truth; the p-value of the Spearman's ρ; the model $R^2$; the mean absolute error (MAE). The test performance is slightly reduced, testifying to poorer generalization as a result of incorporation of some family-specific features.

**(c)** A scatter plot of the incubation periods of the CoV training set compared to the model predictions when trained using family-specific features only.

**(d)** A scatter plot of the incubation periods of the test set compared to the model predictions, when trained using family-specific features only. The box in the bottom right corner contains: Spearman's ρ between predictions and truth; the p-value of the Spearman's ρ; the model $R^2$; the mean absolute error (MAE). The test performance is substantially reduced, testifying to poor generalization and performance of the model trained with family-specific features.

**(e)** A bar plot of the model performance metrics using the family-generic features and an addition of one family-specific feature at each time (X-axes). Of the 4 family-specific features, 3 are zeroed when incorporated to the family-generic model, while the remaining family-specific feature (position change mean) slightly reduces the test performance when included.

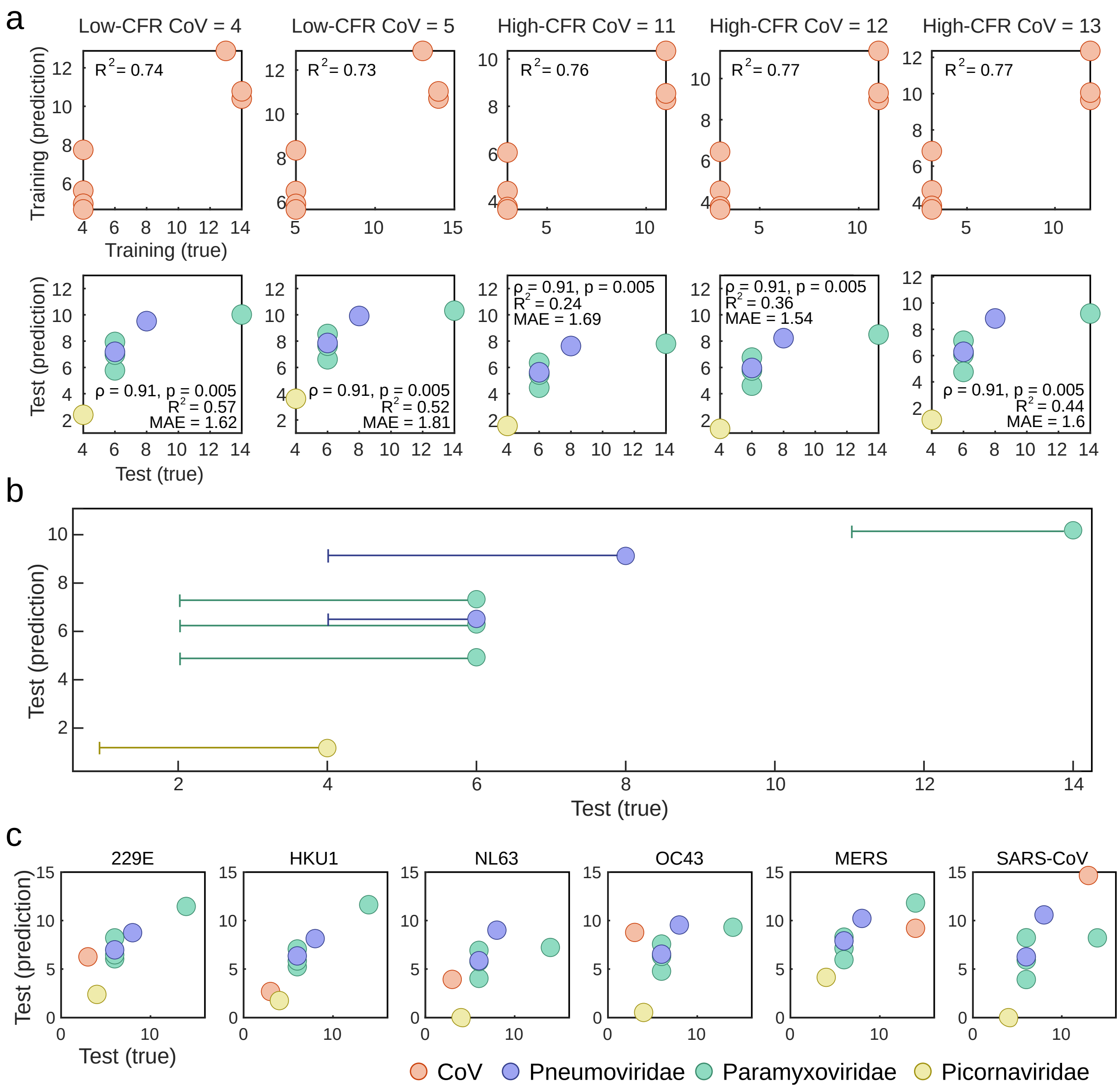


**Supplementary Figure 2. Model performance for ranging incubation time assignment.**

**(a)** Scatter plots of the incubation periods compared to the model predictions across different assigned values of incubation periods. The upper panels show the CoV training set compared to the model predictions when setting different values to the incubation times of CoV (within the literature reported range) in the training set, with the model $R^2$ in the upper left corner. The bottom panels show the incubation periods of the test set compared to the model predictions when setting different values to the incubation times of CoV (within the literature reported range) in the training set. The values in the bottom right or top left corners include: Spearman's ρ between predictions and truth; the p-value of the Spearman's ρ; the model $R^2$; the mean absolute error (MAE). Low-CFR CoV are 229E-CoV, HKU1-CoV, NL63-CoV and OC43-Cov. High CFR-CoV are MERS-CoV, SARS-CoV, and SARS-CoV-2.

**(b)** A scatter plot of the incubation periods of the test set compared to the model predictions, with the full range of incubation period reported in the literature marked for each point on the X-axis (true incubation periods). The incubation period used in the manuscript is marked by a circle and is always the upper limit of the range reported.

**(c)** Scatter plots of the incubation periods of the test set compared to the model predictions when each CoV (other than SARS-CoV-2) is left out of training.

| name | Virus family | Incubation period, upper limit (days) | Number of genes | Genome length, nt | Reference genome accession number | RNA strand polarity |
|---|---|---|---|---|---|---|
| HCoV-229E | *Coronaviridae* | 3 | 7 | 27317 | NC_002645 | + |
| HCoV-NL63 | *Coronaviridae* | 3 | 6 | 27553 | NC_005831 | + |
| HCoV-OC43 | *Coronaviridae* | 3 | 8 | 30741 | NC_006213 | + |
| HCoV-HKU1 | *Coronaviridae* | 3 | 8 | 29926 | NC_006577 | + |
| SARS-CoV | *Coronaviridae* | 13 | 13 | 29751 | NC_004718 | + |
| SARS-CoV-2 | *Coronaviridae* | 14 | 11 | 29903 | NC_045512 | + |
| MERS-CoV | *Coronaviridae* | 14 | 10 | 30119 | NC_019843 | + |
| Measles Virus | *Paramyxoviridae* | 14 | 6 | 15894 | NC_001498 | - |
| Human Parainfluenza virus 1 | *Paramyxoviridae* | 6 | 10 | 15600 | NC_003461 | - |
| Human Parainfluenza virus 2 | *Paramyxoviridae* | 6 | 6 | 15646 | NC_003443 | - |
| Human Parainfluenza virus 3 | *Paramyxoviridae* | 6 | 8 | 15462 | NC_001796 | - |
| Rhinovirus | *Picornaviridae* | 4 | 1 | 7137 | NC_038311 | + |
| Human Metapneumovirus | *Pneumoviridae* | 6 | 8 | 13350 | NC_039199 | - |
| Respiratory Syncytial Virus | *Pneumoviridae* | 8 | 12 | 15191 | NC_001803 | - |

| name | Virus family | CAI | GC content | Position change mean | Position change variance | Mutation rate |
|---|---|---|---|---|---|---|
| HCoV-229E | *Coronaviridae* | 0.66314 | 38.26189 | 0.002825 | 0.000112 | 0.001622 |
| HCoV-NL63 | *Coronaviridae* | 0.63743 | 34.46086 | 0.001657 | 2.98E-05 | 0.001732 |

| | | | | | | |
|---|---|---|---|---|---|---|
| HCoV-OC43 | *Coronaviridae* | 0.65990 | 36.788 | 0.000783 | 4.52E-06 | 0.000857 |
| HCoV-HKU1 | *Coronaviridae* | 0.63454 | 32.05908 | 0.003325 | 8.30E-05 | 0.00206 |
| SARS-CoV | *Coronaviridae* | 0.67680 | 40.76166 | 0.00126 | 4.30E-06 | 0.001489 |
| SARS-CoV-2 | *Coronaviridae* | 0.65464 | 37.97278 | 0.00016 | 2.34E-06 | 7.81E-05 |
| MERS-CoV | *Coronaviridae* | 0.67206 | 41.23643 | 0.001968 | 5.66E-06 | 0.002302 |
| Measles Virus | *Paramyxoviridae* | 0.70395 | 47.4267 | 0.002105 | 7.31E-06 | 0.002627 |
| Human Parainfluenza virus 1 | *Paramyxoviridae* | 0.65739 | 37.24359 | 0.003944 | 8.61E-05 | 0.004174 |
| Human Parainfluenza virus 2 | *Paramyxoviridae* | 0.66245 | 38.40598 | 0.00392 | 6.39E-05 | 0.004534 |
| Human Parainfluenza virus 3 | *Paramyxoviridae* | 0.66421 | 34.52335 | 0.003311 | 3.77E-06 | 0.006644 |
| Rhinovirus | *Picornaviridae* | 0.66992 | 37.4387 | 0.007862 | 2.72E-05 | 0.027469 |
| Human Metapneumovirus | *Pneumoviridae* | 0.66138 | 36.29213 | 0.004148 | 2.61E-05 | 0.006782 |
| Respiratory Syncytial Virus | *Pneumoviridae* | 0.65503 | 33.20387 | 0.000902 | 4.12E-07 | 0.002576 |

**Supplementary Table 1.** The family-generic features for the 14 viruses studied, including viral family, upper limit of the incubation period, genome length, accession number of the reference genome, RNA strand polarity, viral family, codon adaptation index, GC content, mean and variance of changes in positions of the alignment of the virus, and mutation rate.
